# Association of CRP and synovial fluid HMGB1 with Pain in Oligoarticular and Polyarticular Juvenile Idiopathic Arthritis: a cross-sectional study

**DOI:** 10.64898/2026.03.13.711592

**Authors:** Xingzhao Wen, Jacob Rösmark, Anglique Versteegen, Erik Sunderberg, Maria Altman, Cecilia Aulin, Helena Erlandsson Harris

**Author notes:** Correspondence to: Prof. Helena Erlandsson Harris, Center for Molecular Medicine, division of Rheumatology, Department of Medicine Solna, Karolinska Institutet and Karolinska University Hospital, SE-171 76 Stockholm.

## Abstract

**Background:** Pain is one of the most prevalent and distressing symptoms in juvenile idiopathic arthritis (JIA) and often persists despite treatment. Damage-associated molecular patterns (DAMPs), such as high mobility group box 1 (HMGB1) and S100A8/A9, have been implicated in inflammatory activation and nociceptive sensitization, but their associations with pain are not fully characterized in JIA.

**Methods:** Plasma and paired synovial fluid (SF) samples were obtained from patients with oligoarticular and polyarticular JIA from the Juvenile Arthritis Biobank (JABBA). A discovery cohort (n = 79) was used to investigate associations between biomarkers and pain, and these associations were subsequently examined in a validation cohort (n = 38). Levels of HMGB1, S100A8/A9, IL-6, IL-8, C2C, and TRAP5b were measured using ELISA. Associations between biomarkers and patient-reported pain scores were assessed using multivariable linear regression analyses.

**Results:** Plasma and SF levels of most biomarkers did not show significant correlations, except for TRAP5b, which demonstrated a moderate correlation. In the discovery cohort, as multivariable linear regression analyses, both CRP and SF HMGB1 (β = 1.14, 95% CI: 0.21–2.08; β = 1.54, 95% CI: 0.06–3.01 respectively in fully adjusted model) were independently associated with higher pain scores. SF S100A8/A9 (β = 1.00, 95% CI: 0.10–1.89) was additionally associated with pain in fully adjusted models. Sensitivity analyses confirmed the robustness of these findings. These associations were further supported in the validation cohort.

**Conclusions:** Pain in JIA is associated with both systemic CRP and local alarmin markers, with SF HMGB1 showing a particularly robust association. These findings highlight the importance of local joint HMGB1 in pain mechanisms and suggest a potential role for DAMP-mediated pathways in persistent pain in JIA.

## 1. Introduction

Juvenile idiopathic arthritis (JIA) represents the most prevalent rheumatic condition in children before 16 years old [1], affecting 12.8-31.7 out of every 100,000 children each year in Nordic countries [2-4]. JIA is characterized by persistent inflammation in the joints and is associated with pain, stiffness, and structural joint damage [1]. It encompasses seven defined subtypes based on clinical symptoms and laboratory criteria [5].

Despite recent developments regarding treatments, JIA still has a significant impact on the quality of life for patients. Pain is widely recognized as the most frequent and distressing symptom experienced by children with JIA [6-8]. In a cohort of children with polyarticular arthritis, 76% reported pain on more than 60% of days, even while receiving methotrexate, and biologics treatment [8]. Given the high prevalence of persistent pain in JIA despite anti-inflammatory treatment, there is a pressing need to better understand the biological factors that are associated with pain.

Multiple biological pathways may contribute to the pain experience in JIA [9-11]. Among the molecular mediators potentially involved in persistent pain in JIA, alarmins represent a potentially relevant group [12-16]. High mobility group box 1 (HMGB1), a damage-associated molecular pattern (DAMP) released during cellular stress and tissue injury [17], has emerged as a potential mediator of both inflammatory activation and nociceptive sensitization [18, 19]. S100A8/A9, another well-characterized alarmin, was also reported to be involving in pain regulation [15, 20]. However, their association with pain, particularly in JIA and across both circulating and joint-local levels, remains insufficiently investigated.

In this study, we measured HMGB1 and S100A8/A9 in plasma and synovial fluid samples from patients with oligoarticular and polyarticular JIA to investigate their association with pain. In addition, we included inflammatory cytokines (IL-6 and IL-8), tissue-derived mediators reflecting cartilage degradation (C2C) [21] and bone resorption (TRAP5b) [22], as well as circulating CRP, to capture distinct biological pathways potentially linked to pain. Our aim was to identify biomarkers associated with pain in JIA and to provide molecular insights into mechanisms underlying persistent pain in affected children.

## 2. Methods and Materials

Samples from patients with JIA were collected in accordance with the Declaration of Helsinki. Ethical approval was obtained from the North Ethical Committee in Stockholm, Sweden (Dnrs 2009-1139-30-4 and 2010-165-31-2 for Juvenile Arthritis BioBank Astrid Lindgren’s hospital (JABBA)).

### 2.1 Clinical study population and sample collection

Plasma and synovial fluid samples from patients with juvenile idiopathic arthritis (JIA) were obtained from the JIA Biobank (JABBA) at Astrid Lindgren’s Children’s Hospital, Stockholm, Sweden. Clinical and laboratory data collected at the time of sampling were retrieved from medical records and from the Swedish Pediatric Rheumatology Quality Register (Svenska Barnreumaregistret, Omda®). After collection, the samples were processed within four hours of retrieval and stored at - 80 °C freezers prior to analyses.

Patients with oligoarticular and polyarticular JIA who had paired plasma and synovial fluid samples were identified from the registry. For patients with multiple sampling occasions with paired samples, only the earliest paired samples were included in the analyses. Biomarker measurements were performed during two distinct time periods. As a result, biomarker analyses were conducted in separate experimental batches using different commercial assay kits. Because these differences precluded direct pooling of the data, patients analyzed in the later period (n = 79) were used as a discovery cohort to investigate associations between plasma and synovial fluid biomarkers and pain. Patients analyzed in the earlier period (n = 38) were analyzed independently and used as a validation cohort to confirm these associations. A flowchart describing the inclusion steps is shown in Figure 1.

**Figure 1.**
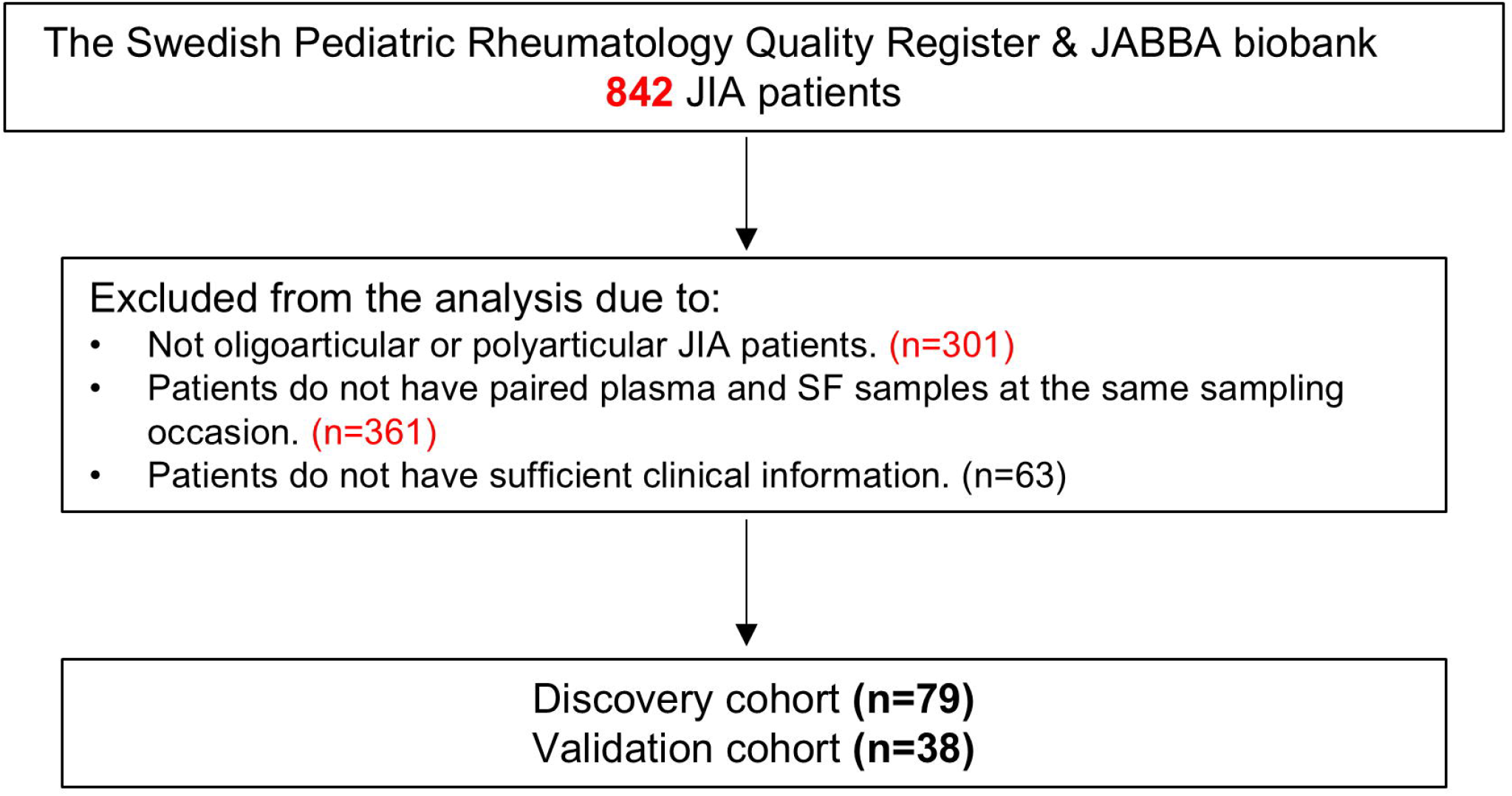
Flow chart illustrating patient recruitment in the current study.

The following clinical data were extracted from the register: date of birth, sex, disease duration (from disease onset to sampling date), number of active joints at visit, CRP and ESR values, presence of autoantibodies (anti-rheumatoid factor (RF) and anti-nuclear antibody (ANA)), treatment, Patient’s Own Reported (PER) Scores of Pain, PER scores of well-being, and the scores of Childhood Health Assessment Questionnaire (CHAQ). Demographic and clinical characteristics of the two cohorts are summarized in Table 1.

### 2.2 Enzyme-linked immunosorbent assays (ELISA)

Plasma and SF samples were thawed only once prior to ELISA assay. SF samples were diluted 1:2 with PBS before analysis.

HMGB1 concentrations were measured using two commercial ELISA kits. Samples from the validation cohort were analyzed using the ST51011 HMGB1 ELISA (IBL International, Germany), whereas samples from the discovery cohort were analyzed using the HMGB1 Express ELISA, an updated assay from IBL International. S100A8/A9 concentrations were measured using different immunoassays in the two cohorts: the Gentian Calprotectin Immunoassay (GCAL®) was used for the validation cohort, and an ELISA kit from R&D Systems (Minneapolis, MN, USA) was used for the discovery cohort. All other biomarkers were measured using the same assays. Levels of IL-6 and IL-8 were determined using ELISA kits from R&D Systems (Minneapolis, MN, USA). The cartilage degradation marker C2C was measured using an ELISA kit from IBEX Pharmaceuticals Inc. (Montréal, Québec, Canada), and the bone resorption marker TRAP5b was quantified using an ELISA kit from Quidel Corporation, USA. All ELISA assays were performed according to the manufacturers’ protocols. Optical density (OD) values were measured using a SpectraMax microplate reader (Molecular Devices, USA). Values below the limit of detection were imputed as half of the lowest detectable concentration.

### 2.3 Statistical Analysis

All statistical analyses were performed using GraphPad Prism (version 9) and R software. Analyses were primarily conducted in the discovery cohort. Prior to analysis, biomarker concentrations measured in plasma and SF were log10-transformed to reduce skewness and approximate a more symmetric distribution. Spearman rank correlation analysis was used to assess correlations between biomarker levels in plasma and SF, as well as correlations between biomarker levels and pain scores. Multivariable linear regression analyses were also performed to evaluate the associations between plasma and SF biomarkers and pain. In unadjusted models, pain score was entered as the dependent variable and each biomarker was analyzed individually as the independent variable. Adjusted models were then constructed to account for potential confounders, including age, sex, disease duration, and treatment status. Treatment status was included as a categorical variable with three levels and modeled using dummy variables. As a sensitivity analysis, patients were dichotomized into low- and high-pain groups based on the median pain score. Least absolute shrinkage and selection operator (LASSO) regression was applied for variable selection to identify candidate biomarkers related to pain. Biomarkers selected by LASSO were subsequently entered into multivariable logistic regression models to assess their associations with pain status, with adjustment for age, sex, disease duration, and treatment status. Spearman correlation and linear regression analyses were then performed to validate the associations in the validation cohort.

## 3. Results

Table 1 presented the demographics and disease characteristics in the two cohorts.

### 3.1 Correlations of the biomarkers between plasma and SF levels

We first performed Spearman rank correlation analyses to examine the associations between plasma and synovial fluid (SF) levels of the selected proteins. Spearman correlation analyses revealed weak or no correlations between plasma and SF levels for most biomarkers, including HMGB1 and S100A8/A9 (Table S1). In contrast, TRAP5b showed a moderate positive correlation between plasma and SF levels (Rho = 0.55, p < 0.001). These findings suggest that, for most markers, local joint changes are not adequately reflected by circulating levels in patients with oligoarticular and polyarticular JIA.

### 3.2 Correlations between plasma/SF biomarkers and pain scores

To preliminarily assess the associations between selected plasma and SF biomarkers and pain, Spearman rank correlation analyses were conducted between biomarker levels and VAS pain scores. Plasma CRP showed a moderate correlation with pain (Rho = 0.281, P = 0.012). SF HMGB1 was also moderately correlated with pain (Rho = 0.268, P = 0.017). SF S100A8/A9 showed a weak correlation that did not reach statistical significance (Rho = 0.221, P = 0.0504). No significant correlations were observed for the other biomarkers in plasma and SF. Detailed results are provided in Table S2.

### 3.3 Multivariable linear regression analysis of biomarkers associated with pain

To account for potential confounding factors, multiple linear regression analyses were performed to examine the associations between biomarkers and pain scores. In the unadjusted analysis, consistent with the Spearman correlation results, only CRP and synovial fluid (SF) HMGB1 were positively associated with pain scores (CRP: β = 1.066, 95% CI: 0.136–1.997; SF HMGB1: β = 1.514, 95% CI: 0.161–2.866). After adjustment for age and sex, the associations between CRP and pain, as well as between SF HMGB1 and pain, remained significant (CRP: β = 1.105, 95% CI: 0.160– 2.050; SF HMGB1: β = 1.524, 95% CI: 0.050–2.999). Further adjustment for disease duration and treatment status yielded similar results. CRP (β = 1.143, 95% CI: 0.208– 2.079) and SF HMGB1 (β = 1.535, 95% CI: 0.061–3.011) remained positively associated with pain scores. In addition, SF S100A8/A9 was associated with increased pain (β = 0.996, 95% CI: 0.100–1.893). Detailed results are shown in Figure 2 and Table 2.

**Figure 2.**
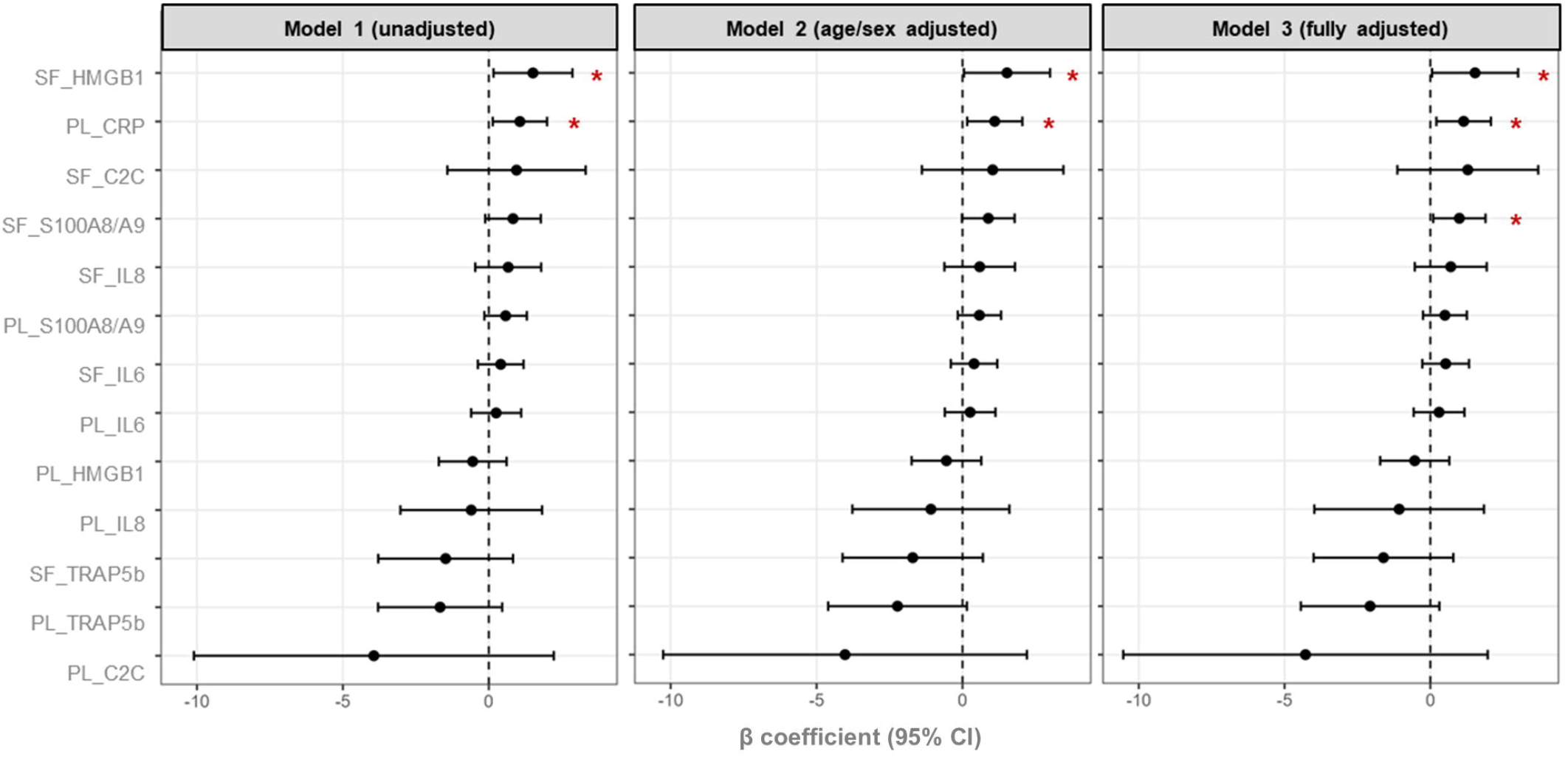
Forest plots showing the associations between biomarkers and pain across three linear regression models. Regression coefficients (β) and corresponding 95% confidence intervals (CI) for each biomarker are displayed in the figure. Model 1: an unadjusted model; Model 2: a model adjusted for age and sex; Model 3: a fully adjusted model including age, sex, disease duration, and treatment. *: P < 0.05.

### 3.4 Sensitivity analysis confirms the robustness of biomarker–pain associations

As a sensitivity analysis, pain was dichotomized into low- and high-pain groups based on the median pain score (Figure 3A), and multivariable logistic regression analyses were performed. LASSO regression with cross-validation was applied to select relevant predictors of high pain from 13 biomarkers. LASSO identified four biomarkers, plasma CRP, SF HMGB1, plasma S100A8/A9, and plasma TRAP5b, as variables with non-zero coefficients (Figure 3B-3C).

**Figure 3.**
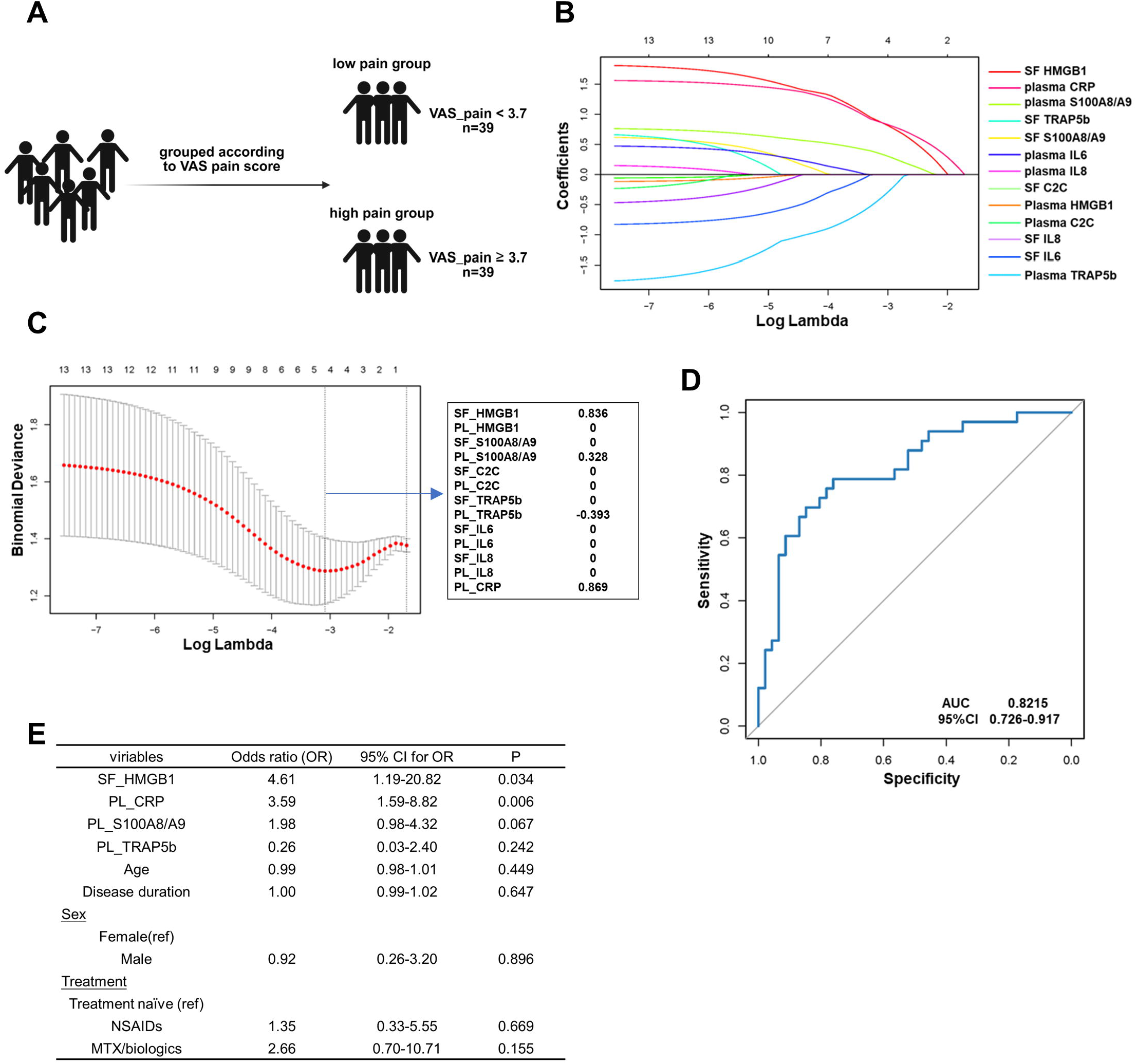
Sensitivity analysis of biomarker–pain associations using multivariable logistic regression. **A)** Schematic diagram of the grouping strategy based on pain median in the cohort. **B-C)** LASSO regression with cross-validation was used to select relevant predictors of high pain from 13 biomarkers, identifying plasma CRP, SF HMGB1, plasma S100A8/A9, and plasma TRAP5b. In **C)**, the red dotted line indicates the cross-validation curve. The dashed vertical lines indicate the optimal values based on the minimum criteria and the 1 − SE criteria. **D)** ROC analysis results of the multivariable logistic regression model, including four selected biomarkers, age, sex, disease duration, and treatment status demonstrating good discrimination between high and low pain groups. **E)** detailed results of multivariable logistic regression model.

These four proteins picked out, together with age, sex, disease duration, and treatment status, were subsequently included in the multivariable logistic regression model. The final model demonstrated good discrimination between patients with high and low pain (Figure 3D). In this model, higher levels of CRP and SF HMGB1 were associated with increased odds of high pain (plasma CRP: OR 3.59, 95% CI 1.59-8.82; SF HMGB1: OR 4.61, 95% CI 1.19-20.82; Figure 3E). Overall, the findings from the logistic regression analysis were consistent with those obtained from the multivariable linear regression models.

### 3.5 Validation the association between CRP/SF HMGB1 and pain in validation cohort

To validate our findings, we investigated the association between plasma CRP, synovial fluid (SF) HMGB1, and pain in a validation cohort (n=38). Spearman correlation analysis showed moderate positive correlations between plasma CRP and SF HMGB1 with pain (CRP: Rho=0.332, P=0.042; HMGB1: Rho=0.358, P=0.028). Subsequent linear regression analysis further confirmed the association between plasma CRP/SF HMGB1 and pain, aligning with the results observed in the discovery cohort (Figure 4B-4C). Taken together, these findings support the robustness and reproducibility of the observed associations.

**Figure 4.**
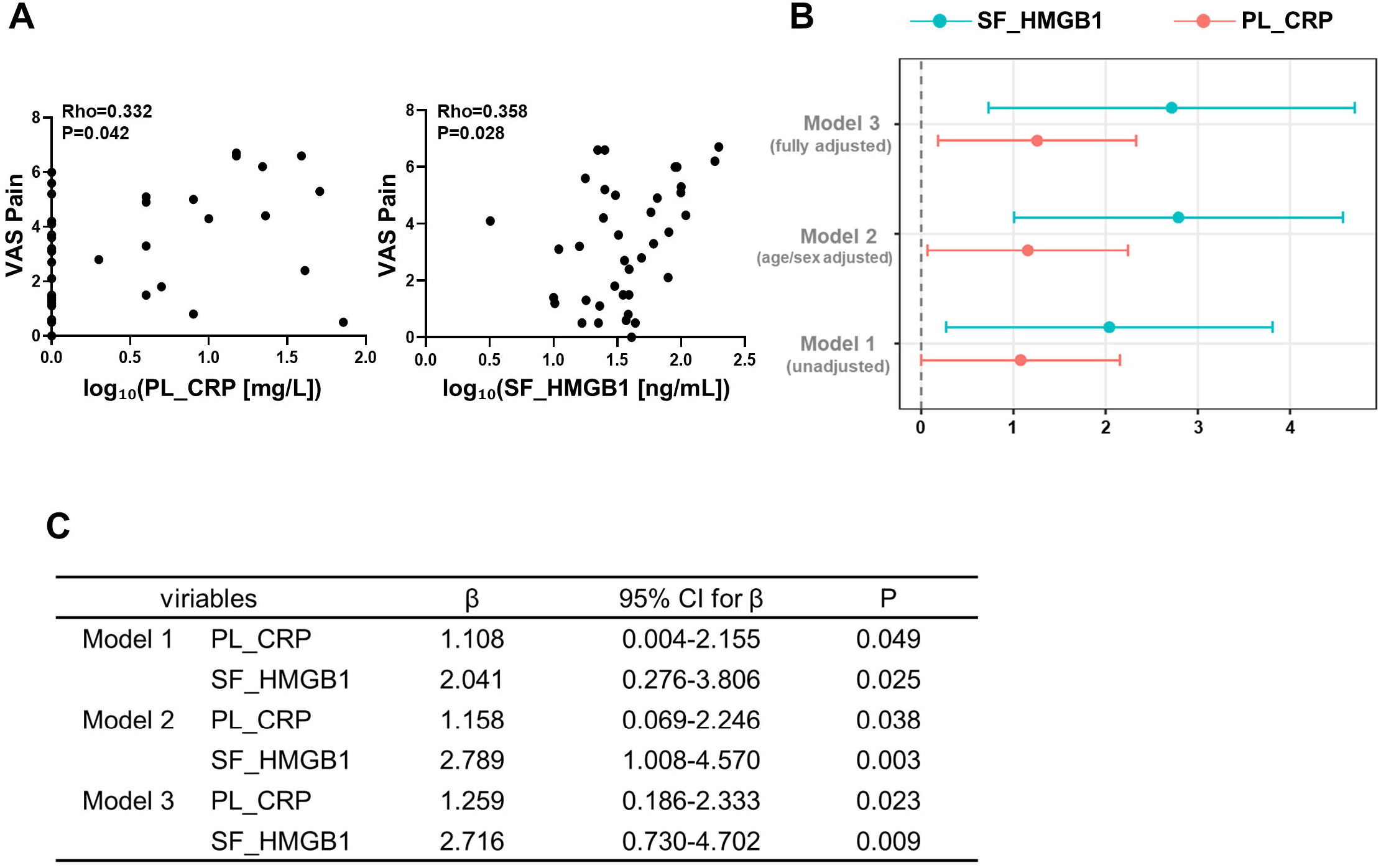
Validation the association between CRP/SF HMGB1 and pain in validation cohort. **A)** scatter plots presenting the correlations of plasma CRP/SF HMGB1 (log10 transformed) with pain in validation cohort. **B)** Forest plots showing the associations between plasma CRP/SF HMGB1 and pain across three linear regression models. Regression coefficients (β) and corresponding 95% CI for plasma CRP and SF HMGB1 are displayed in the figure. Model 1: an unadjusted model; Model 2: a model adjusted for age and sex; Model 3: a fully adjusted model including age, sex, disease duration, and treatment. **C)** detailed results of three linear regression models.

## 4. Discussion

In this study, we investigated the associations between pain and a panel of inflammatory and tissue-derived biomarkers in oligoarticular and polyarticular JIA. We found that pain was associated with synovial fluid HMGB1, as well as with systemic CRP. These associations were consistent across multiple analytical approaches and supported in a validation cohort, highlighting their potential importance in understanding pain in JIA.

Accumulating evidence suggests that DAMPs play an important role in linking tissue inflammation to pain sensitization. HMGB1, a prototypical DAMP released by stressed or damaged cells, has been shown to promote nociceptor sensitization through activation of pattern recognition receptors such as TLR4 and RAGE expressed on sensory neurons, as well as through amplification of local inflammatory responses[16, 19, 23, 24]. In the inflamed synovium, elevated HMGB1 may therefore contribute to pain not only as a marker of tissue damage, but also as an active mediator of peripheral sensitization[24, 25]. S100A8/A9, have also been implicated in inflammatory signaling and may influence pain in arthritis[20, 26-28]. It was reported S100A8/A9 released by synovium cells in inflammatory environment, contributing to inflammatory pain[15]. Despite substantial evidence supporting the role of DAMPs in pain induction, their contribution to pain in patients with JIA has been relatively underexplored. Interestingly, in our previous work, we reported elevated HMGB1 levels in JIA patients and an association with early disease onset [29], suggesting a potential role for HMGB1 in disease pathophysiology. Extending these findings, the present study demonstrates that synovial fluid HMGB1 is independently and consistently associated with pain in patients with oligoarticular and polyarticular JIA across multiple analytical approaches. In contrast, synovial fluid S100A8/A9 was associated with pain only after adjustment for clinical confounders, and this relationship appeared to be influenced by treatment status. These findings indicate that, while both HMGB1 and S100A8/A9 may contribute to pain in JIA, HMGB1 represents a more robust and treatment-independent mediator of pain, whereas the association between S100A8/A9 and pain appears to be more context-dependent. Importantly, our observational findings support the biological plausibility of HMGB1 as a contributor to pain in JIA, as suggested by prior experimental studies on nociceptive signaling. Collectively, these data suggest that HMGB1 may represent a promising therapeutic target in JIA, with the potential to modulate both joint inflammation and pain.

C-reactive protein (CRP), a widely used marker of systemic inflammatory activity, has been associated with pain severity in a range of inflammatory and non-inflammatory conditions, suggesting a link between systemic inflammation and pain[30-32]. In the present study, we observed an association between systemic CRP levels and pain, supporting a role for systemic inflammatory activity in shaping pain experiences in patients with oligoarticular and polyarticular JIA. However, we noticed that CRP levels were very low in a substantial proportion of patients in both cohorts (36 of 79 patients with CRP ≤ 1 mg/L in the discovery cohort and 21 of 38 patients in the validation cohort), indicating that CRP primarily reflects pain-related processes in patients with higher levels of systemic inflammation. In addition, as oligoarticular and polyarticular JIA are characterized by inflammation that is primarily localized to the joints[33, 34], systemic CRP may not accurately capture dynamic changes within the joint. Consistent with this notion, plasma levels of most biomarkers did not show significant correlation with their corresponding synovial fluid concentrations in our analysis. Taken together, our findings indicate that the association between systemic CRP and pain in oligoarticular and polyarticular JIA and this association is most evident in patients with elevated CRP levels. Moreover, systemic inflammation and local synovial mechanisms may contribute to pain through partially distinct pathways.

Neither IL-6 nor IL-8 showed a significant association with pain measures in the present study. While this observation differs from some previous reports [35, 36], such discrepancies may be attributable to differences in study populations, disease stage, and treatment exposure. IL-6 and IL-8 are dynamic cytokines with relatively short half-lives and are strongly influenced by systemic factors and ongoing therapy[37, 38], which may limit their ability to reflect pain mechanisms, particularly in clinically heterogeneous cohorts. These factors may contribute to the lack of association observed in the present study and highlight the complexity of linking individual cytokines to pain outcomes in JIA.

Markers of cartilage degeneration and bone absorption, including C2C and TRAP5b, were not significantly associated with pain in our study. This finding suggests that structural tissue degradation and bone resorption may not directly translate into pain perception in oligoarticular and polyarticular JIA, at least in the absence of overt structural damage. These observations further support the notion that pain in JIA is not solely driven by cumulative tissue damage, but also is influenced by active inflammatory and sensitization-related processes[39].

Several limitations of this study should be acknowledged. First, pain assessment in our study relied on patient-reported measures, which may be influenced by subjective and psychosocial factors not captured in the present analysis. Second, although synovial fluid provides valuable insight into local joint processes, biomarker levels may vary between joints and over time, and single-time-point measurements may not fully reflect dynamic changes in inflammatory and pain-related pathways. And longitudinal investigation is much needed for future studies. Finally, our study cohorts are relatively small due to the difficulty of sampling of SF from juvenile patients, which may have reduced analysis power.

In summary, we investigated the associations of different biomarkers in plasma and SF with pain in oligoarticular and polyarticular JIA patients and identified that plasma CRP and SF HMGB1 are positively associated with pain. These findings provide evidence for the important roles of CRP and HMGB1 in pain in JIA. Clinically, these results suggest that measuring both systemic CRP and SF HMGB1 could provide a more comprehensive understanding of pain in JIA patients and help guide personalized treatment strategies. Specifically, targeting local joint-derived mediators such as HMGB1 may hold promise for more effective pain management.

