## Supplementary material for "Association of CRP and synovial fluid HMGB1 with Pain in Oligoarticular and Polyarticular Juvenile Idiopathic Arthritis: a cross-sectional study": Table 1

**Table 1. Demographics and disease characteristics of the participants**

|  | Discovery cohort | Validation cohort |
| --- | --- | --- |
| Sample size(n) | 79 | 38 |
| Sex(F) | 47(59.5%) | 26 (68.4%) |
| Sex(M) | 32 (40.5%) | 12 (31.6%) |
| Age at sampling (Median, Q1-Q3)  **Subtypes**  Oligoarticular  Seropostive polyarthritis  Seronegative polyarthritis  **Laboratory test** | 12.5(7.8-14.8)  58 (73.4%)  3 (3.8%)  18 (22.8%) | 12.0 (7.5-14.8)  33 (86.8%)  1 (2.6%)  4 (10.5%) |
| CRP Measurement(n) | 79 | 38 |
| median, Q1-Q3 | 4 mg/L (1.0-18.0) | 1 mg/L (1.0-8.5) |
| ESR Measurement(n) | 70 | 24 |
| Median, Q1-Q3  **Autoantibodies** | 2.0 mm/h (1.0-11.0) | 6.0 mm/h (3.25-11.5) |
| RF Measurement(n) | 49 | 22 |
| positive | 4 (8.2%) | 1 (4.5%) |
| ANA Measurement(n) | 68 | 33 |
| positive | 32 (47.0%) | 14 (42.4%) |
| **Clinical parameters** |  |  |
| Active joints (median, Q1-Q3) | 1 (1-4) | 1 (1-2) |
| PER assessment of Pain (median, Q1-Q3)  PER assessment of well-being (median, Q1-Q3) | 3.7 (1.4-6.4)  3.9 (1.3-6.1) | 3.3 (1.4-5.1)  3.0 (0.4-5.5) |
| CHAQ (median, Q1-Q3)  **Treatment**  Treatment naive  NSAIDs  MTX/Biologicals | 0.4 (0.0-0.4)  32 (40.5%)  25(31.6%)  22(27.8%) | 0.5 (0.0-0.6)  16 (42.1%)  12 (31.6%)  10 (26.3%) |

**Table 2. The associations between plasma/SF markers and pain using linear regression analysis**

|  | **Model 1 (unadjusted) .** | | | | **Model 2 (sex and age adjusted) .** | | | | **Model 3 (fully adjusted)* .** | | | |
| --- | --- | --- | --- | --- | --- | --- | --- | --- | --- | --- | --- | --- |
| **Markers** | **β** | **95% CI** | **P** | **R^2^** | **β** | **95% CI** | **P** | **R^2^** | **β** | **95% CI** | **P** | **R^2^** |
| SF_HMGB1 | 1.514 | 0.161-2.866 | 0.028 | 0.061 | 1.524 | 0.050-2.999 | 0.043 | 0.070 | 1.535 | 0.061-3.011 | 0.040 | 0.111 |
| PL_CRP | 1.066 | 0.136-1.997 | 0.025 | 0.063 | 1.105 | 0.160-2.050 | 0.022 | 0.075 | 1.143 | 0.208-2.079 | 0.017 | 0.129 |
| SF_C2C | 0.950 | -1.419-3.319 | 0.427 | 0.008 | 1.034 | -1.390-3.458 | 0.398 | 0.017 | 1.285 | -1.130-3.701 | 0.292 | 0.063 |
| SF_S100A8/A9 | 0.831 | -0.122-1.784 | 0.060 | 0.044 | 0.884 | -0.018-1.786 | 0.055 | 0.056 | 0.996 | 0.100-1.893 | 0.030 | 0.108 |
| SF_IL8 | 0.665 | -0.464-1.794 | 0.244 | 0.017 | 0.589 | -0.620-1.798 | 0.335 | 0.020 | 0.701 | -0.532-1.934 | 0.261 | 0.065 |
| PL_S100A8/A9 | 0.577 | -0.150-1.304 | 0.118 | 0.031 | 0.580 | -0.162-1.322 | 0.124 | 0.039 | 0.501 | -0.250-1.252 | 0.188 | 0.071 |
| SF_IL6 | 0.409 | -0.374-1.192 | 0.302 | 0.014 | 0.395 | -0.400-1.191 | 0.325 | 0.021 | 0.527 | -0.274-1.327 | 0.194 | 0.070 |
| PL_IL6 | 0.252 | -0.605-1.109 | 0.560 | 0.004 | 0.262 | -0.606-1.130 | 0.550 | 0.012 | 0.300 | -0.571-1.171 | 0.494 | 0.054 |
| PL_HMGB1 | -0.551 | -1.712-0.610 | 0.348 | 0.011 | -0.550 | -1.744-0.643 | 0.361 | 0.019 | -0.535 | -1.721-0.651 | 0.371 | 0.059 |
| PL_IL8 | -0.602 | -0.303-1.826 | 0.610 | 0.014 | -1.083 | -3.773-1.607 | 0.407 | 0.022 | -1.069 | -3.978-1.840 | 0.446 | 0.085 |
| SF_TRAP5b | -1.478 | -3.788-0.832 | 0.207 | 0.021 | -1.705 | -4.110-0.699 | 0.162 | 0.033 | -1.607 | -4.003-0.789 | 0.186 | 0.071 |
| PL_TRAP5b | -1.668 | -3.794-0.458 | 0.122 | 0.031 | -2.227 | -4.604-0.150 | 0.066 | 0.052 | -2.066 | -4.442-0.311 | 0.0874 | 0.086 |
| PL_C2C | -3.938 | -10.104-2.227 | 0.207 | 0.021 | -4.025 | -10.257-2.208 | 0.202 | 0.029 | -4.282 | -10.533-1.969 | 0.176 | 0.072 |

*: The model adjusted by sex, age, disease duration, and treatment.
