## Supplementary table for "Association of CRP and synovial fluid HMGB1 with Pain in Oligoarticular and Polyarticular Juvenile Idiopathic Arthritis: a cross-sectional study"

**Supplementary tables**

**Supplementary table 1. Correlations of proteins selected between plasma and SF**

| **Proteins** | **Spearmen r** | **P value** |
| --- | --- | --- |
| HMGB1 | 0.016 | 0.888 |
| S100A8/A9 | 0.080 | 0.485 |
| C2C | 0.082 | 0.472 |
| TRAP5b | 0.550 | 1.52E-07 |
| IL-6 | 0.028 | 0.809 |
| IL-8 | 0.240 | 0.294 |

**Supplementary table 2. Correlations between plasma/SF proteins selected and vas pain in patients**

| **Proteins** | **Spearmen r** | **P value** |
| --- | --- | --- |
| Plasma_HMGB1 | -0.040 | 0.725 |
| Plasma_S100A8/A9 | 0.154 | 0.175 |
| Plasma_C2C | -0.106 | 0.350 |
| Plasma_TRAP5b | -0.216 | 0.056 |
| Plasma_IL-6 | 0.016 | 0.891 |
| Plasma_IL-8 | -0.300 | 0.186 |
| Plasma_CRP | 0.281 | 0.012 |
| SF_HMGB1 | 0.268 | 0.017 |
| SF_S100A8/A9 | 0.221 | 0.0504 |
| SF_C2C | 0.112 | 0.327 |
| SF_TRAP5b | -0.144 | 0.206 |
| SF_IL-6 | 0.085 | 0.455 |
| SF_IL-8 | 0.143 | 0.208 |
